## Supplemental Figures for "Cell cycle and Age-Related Modulations of Mouse Chromosome Stiffness"

### Supplement Fig.1

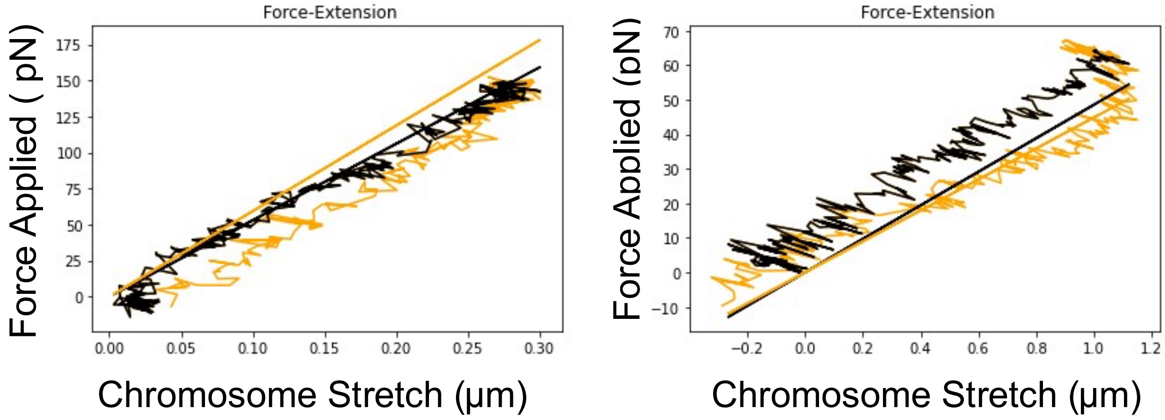


**Supplement Figure 1. Representative images of oocyte chromosome stretching.**

Left panel: MI oocyte chromosome stretching. Right panel: MII oocyte chromosome stretching. The black line (representing the stretching process) and the yellow line (representing the retraction process) almost overlap with each other, which indicates that oocyte chromosomes display elastic properties, and the stretching process is reversible.

### Supplement Fig.2

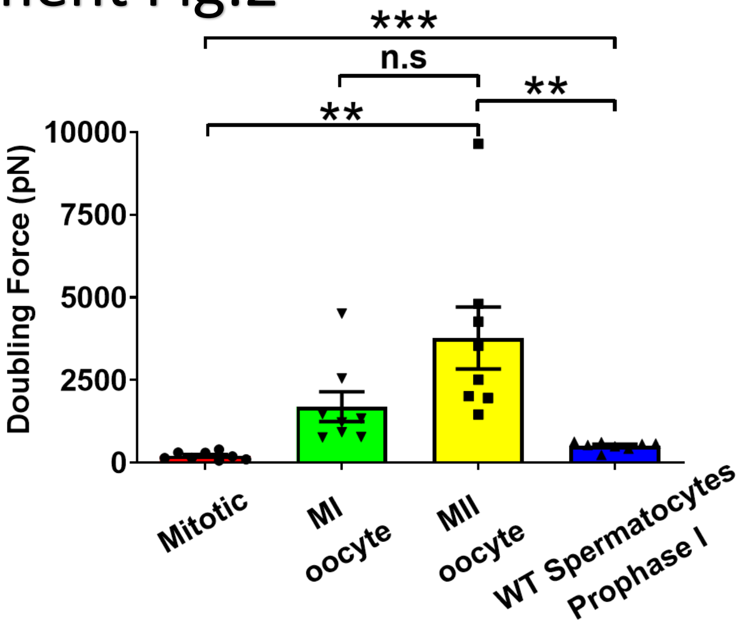


**Supplement Figure 2. Chromosome doubling force comparison across different types of cells.** Chromosome doubling force comparison between mitotic cells (n=8), WT spermatocytes at prophase I (n=8), MI oocytes (n=8) and MII oocytes (n=8).

### Supplement Fig.3

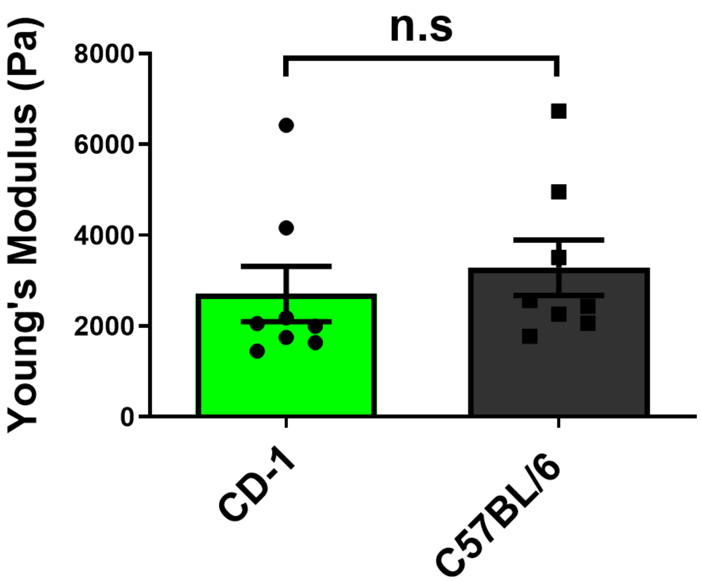


**Supplement Figure 3. Comparison of chromosome stiffness between CD-1 and C57BL/6 mice.**

Young's Modulus of chromosomes was measured and compared between CD-1 mice (n=8) and C57BL/6 mice (n=8). There are no significant stiffness differences between these two mouse lines (2710 ± 610 Pa in CD-1 versus 3290 ± 610 Pa in C57BL6/J, P = 0.512). Data are presented as mean ± SEM, with statistical analysis performed using t-test.

### Supplement Fig.4

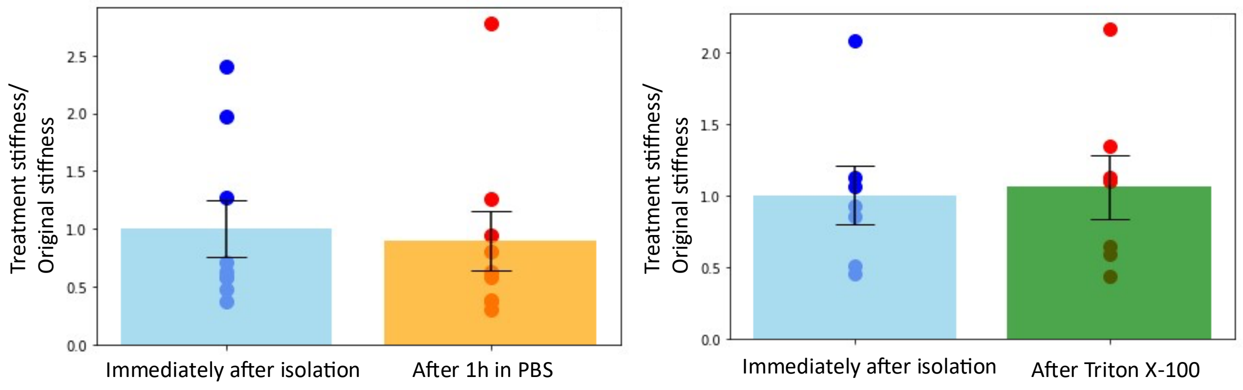


**Supplement Figure 4. Chromosome stiffness is stable in PBS solution and resistant to Triton X-100 treatment.**

Left panel: chromosome stiffness immediately after isolation (1.000 ± 0.2419, n=9) is not significantly different from that sitting in PBS for 1 hour (0.8951 ± 0.2563, n=9, P= 0.7697). Right panel: chromosome stiffness immediately after isolation (1.000 ± 0.2049, n=7) shows no significant change after treatment with 0.05% Triton X-100 for 10 minutes (1.057 ± 0.2226, n=7, P= 0.8527). Data are presented as mean ± SEM, with statistical analysis performed using t-test.
